# Drug-tolerant persister cells reallocate carbon sources to fuel antioxidant metabolism for survival

**DOI:** 10.64898/2026.01.05.697775

**Authors:** Melvin Li, Bradley Priem, Luke V. Loftus, Michael J. Betenbaugh, Kenneth J. Pienta, Sarah R. Amend

## Abstract

Therapy resistance is the leading cause of cancer-related deaths. Drug-tolerant persister cells (DTPs) represent a major barrier to cancer cure, mediating resistance through adaptive cell state transitions and driving tumor progression. Here, we investigate metabolic differences between DTPs and drug-sensitive cancer cells using integrated fluxomics. Proteomic profiling and extracellular flux analyses revealed that DTPs upregulate glycolysis and gluconeogenesis while reducing oxidative phosphorylation, indicating a shift in central carbon metabolism. Isotope tracing and metabolic modeling demonstrate that DTPs utilize glucose to fuel the pentose phosphate pathway (PPP) to generate NADPH and metabolize glutamine to provide carbons for the PPP via gluconeogenesis. Integrating our multi-omic datasets into a genome-scale model identified that DTPs sustain antioxidant metabolism by decreasing fluxes of other NADPH-consuming reactions upon *in silico* PPP knockout. These findings reveal a systems-level shift in DTP metabolism that maintains antioxidant activity for cell survival, highlighting potential new targets and treatment paradigms to overcome therapy resistance.

## INTRODUCTION

Therapy resistance is responsible for 90% of cancer-related deaths and remains a major barrier to cancer cure^1^. Genotoxic chemotherapies, ranging from platinum-based drugs, antimetabolites, and topoisomerase inhibitors, are standard systemic treatments used across all types of solid and blood tumors^2^. While these drugs are initially effective in reducing tumor burden, subpopulations of cancer cells that survive treatment remain – ultimately leading to recurrence and poor prognosis in patients. Classic models of therapy resistance are explained through tumor cell heterogeneity, suggesting that treatment selects for pre-existing clonal populations harboring specific genetic mutations that confer resistance^2^. In recent years, cancer cells have been shown to also adopt non-genetic mechanisms of resistance that are driven by epigenetic and phenotypic plasticity^2^.

Drug-tolerant persister cells (DTPs) emerge in response to chemotherapy through transient and reversible cell state changes^3^. These cells adopt a slow cycling or quiescent phenotype that promotes survival under therapeutic pressure and later seed recurrence after the tumor is no longer clinically detectable^3^. DTPs have been detected across various tumor types (breast, prostate, lung, and melanoma) and induced by many classes of chemotherapy and targeted therapies ^3,4^. Characterizing the phenotypic plasticity of the DTP state will provide a better understanding of resistance mechanisms and reveal potential vulnerabilities to improve disease outcomes. Past studies have shown that DTPs present with stem-like characteristics and altered cell metabolism when compared to their non-persister counterparts^3^.

Cell metabolism is an adaptive process that responds dynamically to nutrient availability, energy demand, and various sources of stress^5,6^. It consists of a complex network linking the epigenome, transcriptome, proteome, metabolome, and fluxome^7–10^. This network is continually shaped by factors such as interactions with the extracellular environment, differential utilization of carbon and nitrogen sources, and rewiring of pathway fluxes^11^. Cancer cells adopt transient metabolic states to allocate resources to grow, proliferate, and resist standard-of-care therapies^12,13^. Depending on the tissue of origin, type of stressors the tumors experience, and disease status, different metabolic pathways can be utilized to fulfill the needs of rapid proliferation and cancer cell survival^14^. Therapy-induced DTPs have been shown to increase fatty acid uptake, fatty acid oxidation and mitochondrial respiration while shifting away from glycolysis^3,15^. Although DTP metabolism has been studied in isolated metabolic pathways, metabolic rewiring at the systems level in DTPs has been largely unexplored.

In this study, we employ multidisciplinary approaches from cancer cell biology, analytical chemistry, and systems biology to investigate the metabolic alterations in the cancer cells that persist after cisplatin treatment. We demonstrate through multi-omic analyses that DTPs increase glucose and glutamine uptake, but rather than shuttling them into ATP-generating pathways, they allocate those carbon sources to generate antioxidants for cell survival. Characterization of the DTP metabolic proteome revealed increase expression of key enzymes involved in glycolysis and gluconeogenesis while decreasing expression of oxidative phosphorylation (OXPHOS)-related proteins. Through isotope tracing and ^13^C-metabolic flux analysis (^13^C-MFA), we quantitatively mapped metabolic flux differences between DTPs and parental cells, showing the rewiring of glucose and glutamine to fuel the oxidative pentose phosphate pathway (oxPPP) in DTPs. We then integrated our proteomic and ^13^C-MFA datasets into a genome scale metabolic model to assess the role of pentose phosphate pathway-derived NADPH on antioxidant metabolism, providing a robust systems-level view on metabolic reprogramming in DTPs surviving cisplatin. *In silico* knockout of the oxPPP in DTPs revealed a possible compensatory mechanism to sustain antioxidant capacity through the reduction of folate metabolism. Our findings provide new insights into the metabolic flexibility of DTPs and highlight potential vulnerabilities to target this transient resistant state.

## RESULTS

### Cells that persist after cisplatin treatment increase glucose metabolism and decrease oxidative phosphorylation

Prostate cancer PC3 parental cells were treated with an LD_50_ dose of cisplatin for 72 hours, treatment was removed, and cultures were maintained to 10 days post-treatment removal. We have previously shown that cells continue to die days after treatment removal, with the death rate plateauing at 10 days^16^. Cells surviving at this timepoint are classified as drug-tolerant persister cells (DTPs) (**Fig. 1A**). The DTPs in our model system present with a non-proliferative phenotype, increased genomic content, and an increase in cell size^16^. We profiled proteomic differences between DTPs and parental cells. Overall, 9,277 proteins were detected. Sample groups (DTPs and parental cells) exhibited distinct proteomic profiles and clustered independently in principal component analysis (PCA) (**Fig. 1B**). Differential expression analysis identified 6,125 differentially expressed proteins between sample groups, with 2,451 enriched in DTPs and 3,674 upregulated proteins in parental cells (**Fig. 1C**). Canonical DTP markers, ALDH1 and CD44, were increased at the protein level in the DTP population when compared to parental cells (**Fig. 1C**)^3^.

**Fig. 1.**
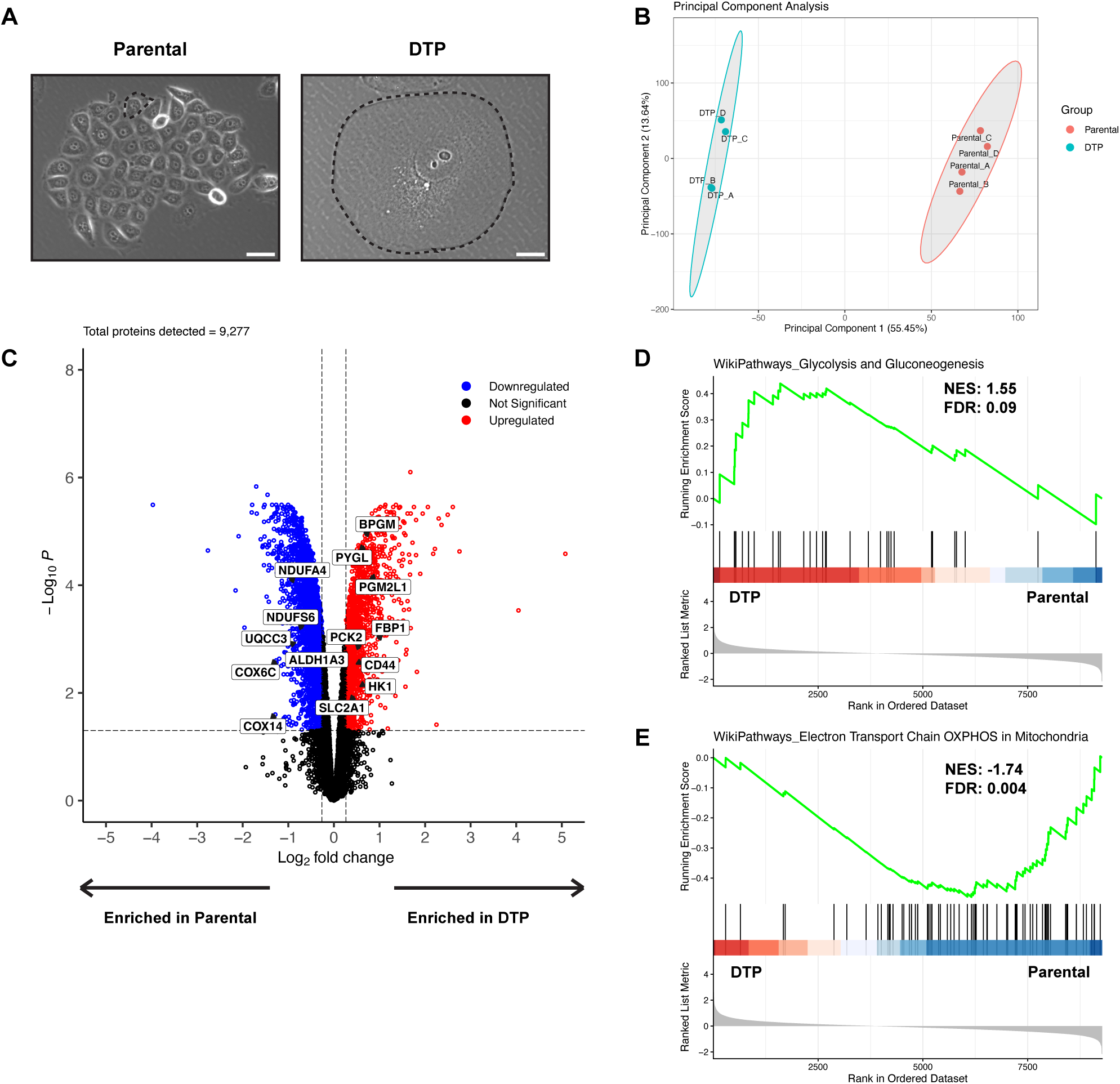
Cells that persist after cisplatin treatment increase glucose metabolism and decrease oxidative phosphorylation. (**A**) Phase contrast images of parental PC3 cells and drug-tolerant persister cells (DTPs) surviving cisplatin treatment. Black dotted lines indicate the cell border of parental cells and DTPs. Scale bar: 100 µm. (**B**) Principal component analysis (PCA) plot of parental cells and DTPs based on log_2_ protein expression. (**C**) Volcano plot of differentially expressed proteins in DTPs vs parental cells. Log_2_ fold change cutoff was 0.263 and adjusted p-value cutoff was 0.05. (**D**) WikiPathways Glycolysis and Gluconeogenesis pathway enrichment analysis between DTPs and parental cells. (**E**) WikiPathways Electron Transport Chain OXPHOS in Mitochondria pathway enrichment analysis between DTPs and parental cells. n = 4 biological replicates for parental cells and DTPs (**B-E**). Adjusted p-values were generated in R using Benjamini-Hochberg correction (**C**). False discovery rates (FDR) and normalized enrichment scores (NES) were generated for pathway enrichment analyses in R. FDR cutoff was set to 0.10 (**D-E**).

It has been previously reported that DTPs in melanoma and acute myeloid leukemia (AML) models increase OXPHOS and shift away from glycolysis, leading to a metabolic vulnerability to Complex I inhibition^15,17,18^. Interestingly in our model system, key proteins mediating glycolysis and gluconeogenesis such as SLC2A1, HK1, and FBP1, were increased in DTPs, while proteins mediating OXPHOS such as COX6C, COX14, and NDUFA4 were decreased (**Fig. 1C**). These findings were further supported by the enrichment of the WikiPathways Glycolysis and Gluconeogenesis gene set and a depletion in the WikiPathways Oxidative Phosphorylation gene set in DTPs via pathway enrichment analysis (**Fig. 1D-E**). Based on these data, we hypothesize that DTPs surviving cisplatin increase glucose uptake and production while shifting mitochondrial metabolism away from ATP production.

### DTPs increase glucose uptake and decrease flux through glycolysis

Given the enrichment of proteins mediating glycolysis and gluconeogenesis in DTPs, we next investigated how glucose is metabolized between DTPs and parental cells. Glucose is imported into the cell and typically metabolized to pyruvate. As described by the Warburg Effect, cancer cells utilize pyruvate to generate lactate, releasing it into the extracellular space at high concentrations^19^. We measured the extracellular fluxes of glucose uptake and lactate excretion and found that DTPs had a 5.9-fold increase in glucose uptake rate compared to parental cells (**Fig. 2A**). Despite the increase in glucose uptake rate, DTPs excreted less lactate over time (**Fig. 2B**). To investigate how glucose is metabolized intracellularly, we cultured cells in media containing [U-^13^C] glucose and dialyzed FBS and analyzed the contribution of the isotope tracer to downstream glycolytic metabolites (**Fig. 2C**). While we observed increased glucose import in DTPs, we found a reduction in [U-^13^C] glucose contribution to glucose 6-phosphate (G6P) when compared to parental cells (**Fig. 2D**). This was perplexing since G6P is the first metabolite downstream of glucose import, where glucose gets phosphorylated at the C6-position^20^. This suggests that there may be another carbon source contributing to the G6P pool. One possible mechanism is through phosphoglucose isomerase (PGI), which can reversibly generate G6P from fructose 6-phosphate (F6P)^21^. We also observed a dramatic drop in contribution of [U-^13^C] glucose to the fructose 1,6-bisphosphate (FBP) pool, suggesting that the tracer is diluted by an unlabeled carbon source (**Fig. 2E**). The contribution of [U-^13^C] glucose to the other downstream glycolytic metabolites was decreased in DTPs when compared to parental cells (**Fig. 2E**). We measured tracer incorporation into extracellular lactate with [1,2-^13^C] glucose using gas chromatography mass spectrometry (GC-MS) and observed decreased levels of m+2 lactate in the media of DTPs. Taken together, these data demonstrate that DTPs decrease glucose contribution to glycolysis and reduce glycolytic flux (**Fig. 2F-G**).

**Fig. 2.**
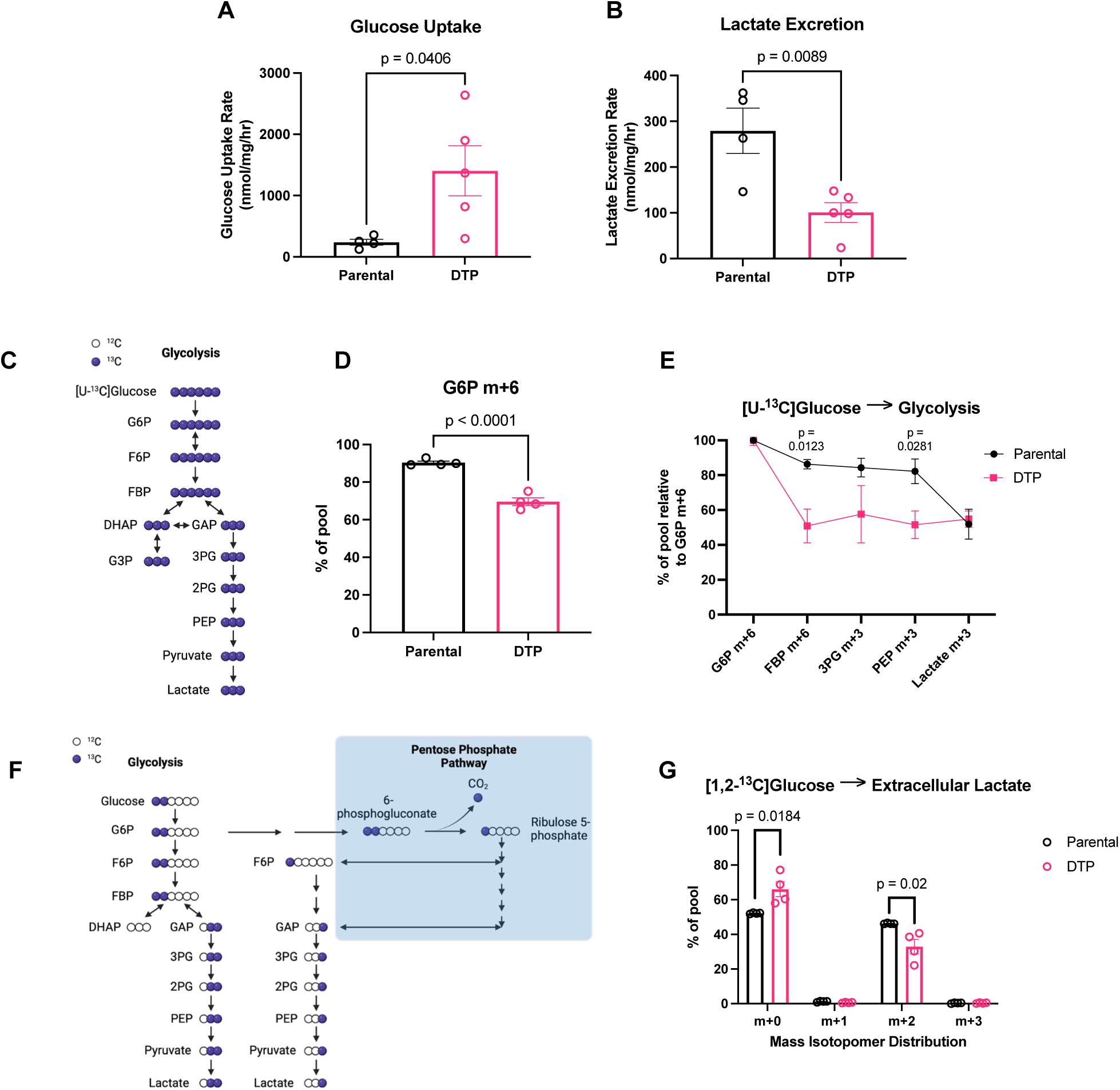
DTPs increase glucose uptake and decrease flux through glycolysis. (**A**) Glucose uptake rate of parental cells and DTPs. (**B**) Lactate excretion rate of parental cells and DTPs. (**C**) Schematic of ^13^C-atom transitions in the glycolytic pathway from [U-^13^C] glucose. Blue circles denote ^13^C. White circles denote ^12^C. Created in BioRender. Li, M. (2026) https://BioRender.com/vwr1a3f. (**D**) Labeling of intracellular G6P m+6 from [U-^13^C] glucose in parental cells and DTPs. (**E**) Labeling of intracellular glycolytic metabolites from [U-^13^C] glucose relative to intracellular G6P m+6 in parental cells and DTPs. (**F**) Schematic of ^13^C-atom transitions in the glycolytic pathway from [1,2-^13^C] glucose. Blue circles denote ^13^C. White circles denote ^12^C. Created in BioRender. Li, M. (2026) https://BioRender.com/hfybwb9. (**G**) Labeling of extracellular lactate m+2 from [1,2-^13^C] glucose from media of parental cells and DTPs. n = 4 biological replicates and n = 5 biological replicates for parental cells and DTPs, respectively (**A-B**); n = 4 biological replicates (**D-E, G**). Data are presented as mean ± s.e.m. P-values were calculated via unpaired Student’s two-tailed t-test and exact values are indicated in the figure. Normal distributions were confirmed for all data with a Shapiro-Wilk test. Abbreviations: glucose 6-phosphate (G6P); fructose 1,6-bisphosphate (FBP); 3-phosphoglycerate (3PG); phosphoenolpyruvate (PEP).

### Glutamine is the major carbon source for the TCA cycle in parental cells and DTPs

Downstream from glycolysis, pyruvate is oxidized to acetyl-CoA and enters the mitochondria and metabolized through the TCA cycle^22^. Through [U-^13^C] glucose tracing, we found that DTPs exhibit decreased labeling of m+2 citrate, suggesting decreased glucose-derived influx into the TCA cycle when compared to parental cells (**Fig. 3A-B**). In both parental cells and DTPs, labeling fractions of metabolites downstream of citrate decreased dramatically to under 10%, indicating that the [U-^13^C] glucose tracer was diluted by an unlabeled carbon source (**Fig. 3B**). The dilution of the [U-^13^C] glucose tracer can be first observed in the labeling of m+2 succinate. Succinate is downstream of alpha-ketoglutarate, which is a key node where glutamine-derived carbons can enter the TCA cycle. It has been shown in various studies that glutamine is another major source of carbon for the TCA cycle in cancer cells ^23–25^, so we hypothesized that both parental cells and DTPs were utilizing glutamine to fuel the TCA cycle instead of glucose.

**Fig. 3.**
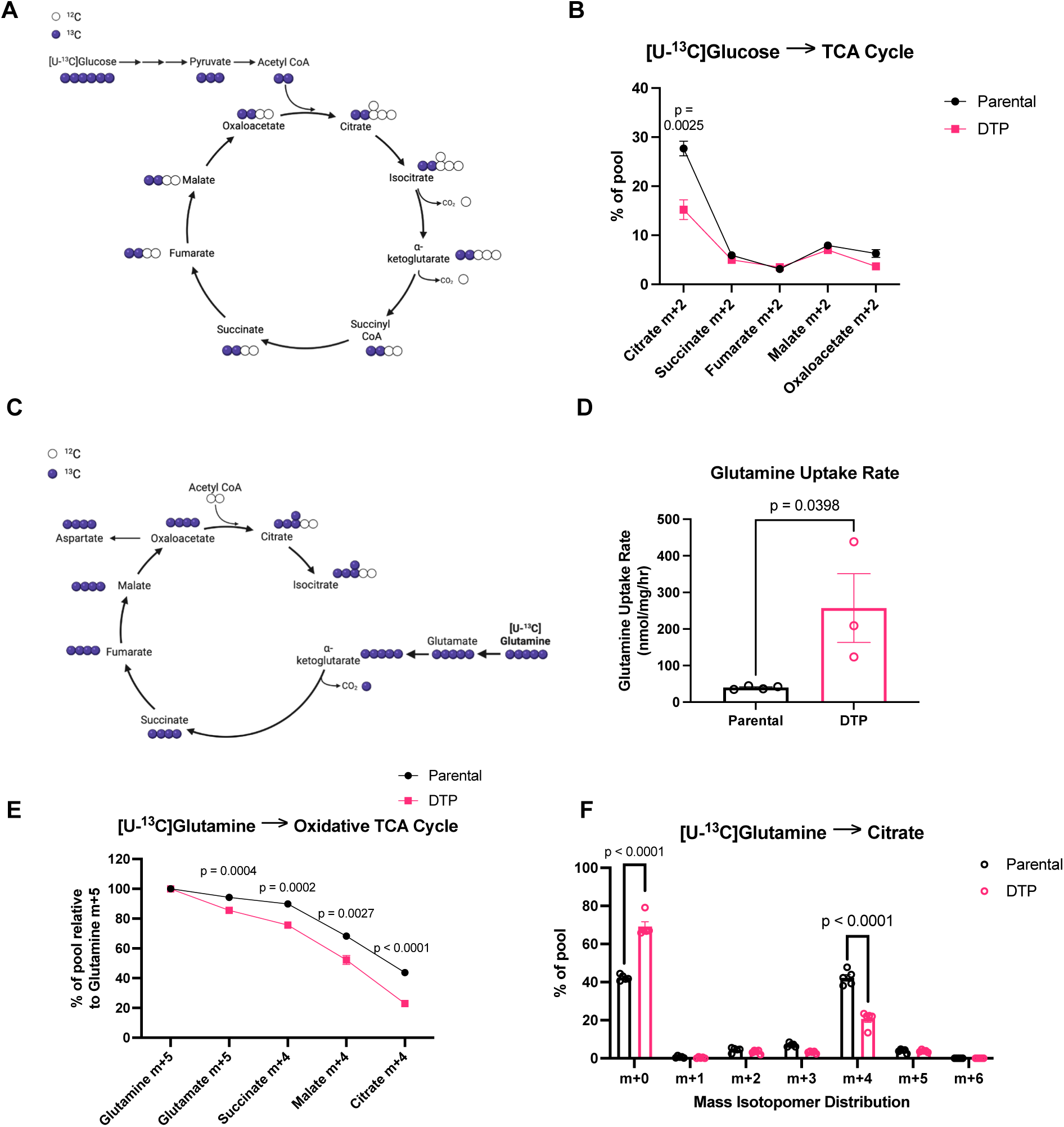
DTPs decrease glutamine contribution into the TCA cycle. (**A**) Schematic of ^13^C-atom transitions in the TCA cycle from [U-^13^C] glucose. Blue circles denote ^13^C. White circles denote ^12^C. Created in BioRender. Li, M. (2026) https://BioRender.com/ufaoeuo. (**B**) Labeling of intracellular TCA cycle metabolites from [U-^13^C] glucose in parental cells and DTPs. (**C**) Schematic of ^13^C-atom transitions in the TCA cycle from [U-^13^C] glutamine. Blue circles denote ^13^C. White circles denote ^12^C. Created in BioRender. Li, M. (2026) https://BioRender.com/ay5843y. (**D**) Glutamine uptake rate of parental cells and DTPs. (**E**) Labeling of intracellular TCA cycle metabolites from [U-^13^C] glutamine relative to intracellular glutamine m+5 in parental cells and DTPs. (**F**) Mass isotopomer distribution (MID) of citrate from [U-^13^C] glutamine in parental cells and DTPs. n = 4 biological replicates (**B**); n = 4 biological replicates and n = 3 biological replicates in parental cells and DTPs, respectively (**D**); n = 5 biological replicates (**E-F**). Data are presented as mean ± s.e.m. P-values were calculated via unpaired Student’s two-tailed t-test and exact values are indicated in the figure. Normal distributions were confirmed for all data with a Shapiro-Wilk test.

### DTPs decrease glutamine contribution into the TCA cycle

We measured the contribution of [U-^13^C] glutamine to TCA cycle metabolites (**Fig 3C**). Carbons from [U-^13^C] glutamine incorporate into glutamate via glutaminase (GLS) activity to generate glutamate m+5, followed by its conversion to alpha-ketoglutarate (*α*-KG) m+5 through glutamate dehydrogenase (GDH) (**Fig 3C**). *α*-KG is then decarboxylated and leads to m+4 labeling of the downstream TCA cycle metabolites such as succinate, fumarate, malate, and oxaloacetate (**Fig 3C**). m+4 citrate is generated as m+4 oxaloacetate reacts with unlabeled acetyl-CoA (**Fig 3C**). Cells were cultured in [U-^13^C] glutamine-containing media and media was sampled every hour for analysis of the extracellular rate of glutamine uptake. DTPs exhibited increased glutamine consumption over time when compared to parental cells (**Fig 3D**). However, we observed that despite the increase in glutamine uptake rate, DTPs show decreased [U-^13^C] glutamine contribution to the intracellular TCA cycle metabolites compared to parental cells (**Fig 3E**). For example, there was decreased enrichment of m+4 citrate from [U-^13^C] glutamine in DTPs, suggesting that other carbon sources are contributing to the citrate pool (**Fig. 3F**). In addition, labeling of m+5 citrate from [U-^13^C] glutamine was lower than 10% in both sample groups, indicating that glutamine does not fuel reductive carboxylation (**Fig. 3F**).

### ^13^C-metabolic flux analysis reveals the reallocation of glucose and glutamine to fuel the pentose phosphate pathway in DTPs

One limitation of solely using mass isotopomer distributions (MIDs) to assess intracellular metabolic activity is the inability to report absolute fluxes of reactions in a pathway. MID data allows us to make conclusions on the relative pathway contributions to the production of a metabolite and therefore can only provide qualitative information^26,27^. To quantify the differences in absolute fluxes of metabolic reactions between parental cells and DTPs, we employed computational modeling to perform ^13^C-metabolic flux analysis (^13^C-MFA) based on established methods by integrating the MID data from the [U-^13^C] glucose and [U-^13^C] glutamine experiments into a defined metabolic model and constraining the model using the measured extracellular fluxes^28,29^.

^13^C-MFA revealed that parental cells primarily metabolized glucose via glycolysis, with glucose-derived pyruvate and lactate production at 0.48 µmol/mg/hr and at 0.28 µmol/mg/hr, respectively (**Fig. 4A**). These fluxes are consistent with other reports of glycolytic flux in proliferating cancer cells^30–32^. In contrast, DTPs metabolized glucose by shuttling G6P into the oxidative pentose phosphate pathway (oxPPP) at a rate of 8.74 µmol/mg/hr, resulting in the production of pentose 5-phosphate (P5P) (**Fig. 4B**). DTPs also exhibit high flux through the non-oxidative pentose phosphate pathway (non-oxPPP), feeding carbons from P5P back into the glycolytic pathway through the generation of fructose 6-phosphate (F6P) and glyceraldehyde 3-phosphate (GAP) (**Fig. 4B**). Interestingly, the net fluxes of the glycolytic reactions in DTPs indicate that glycolysis is operating in the reverse direction, suggesting that in DTPs activate gluconeogenesis with carbons derived from pyruvate (**Fig. 4B**).

**Fig. 4.**
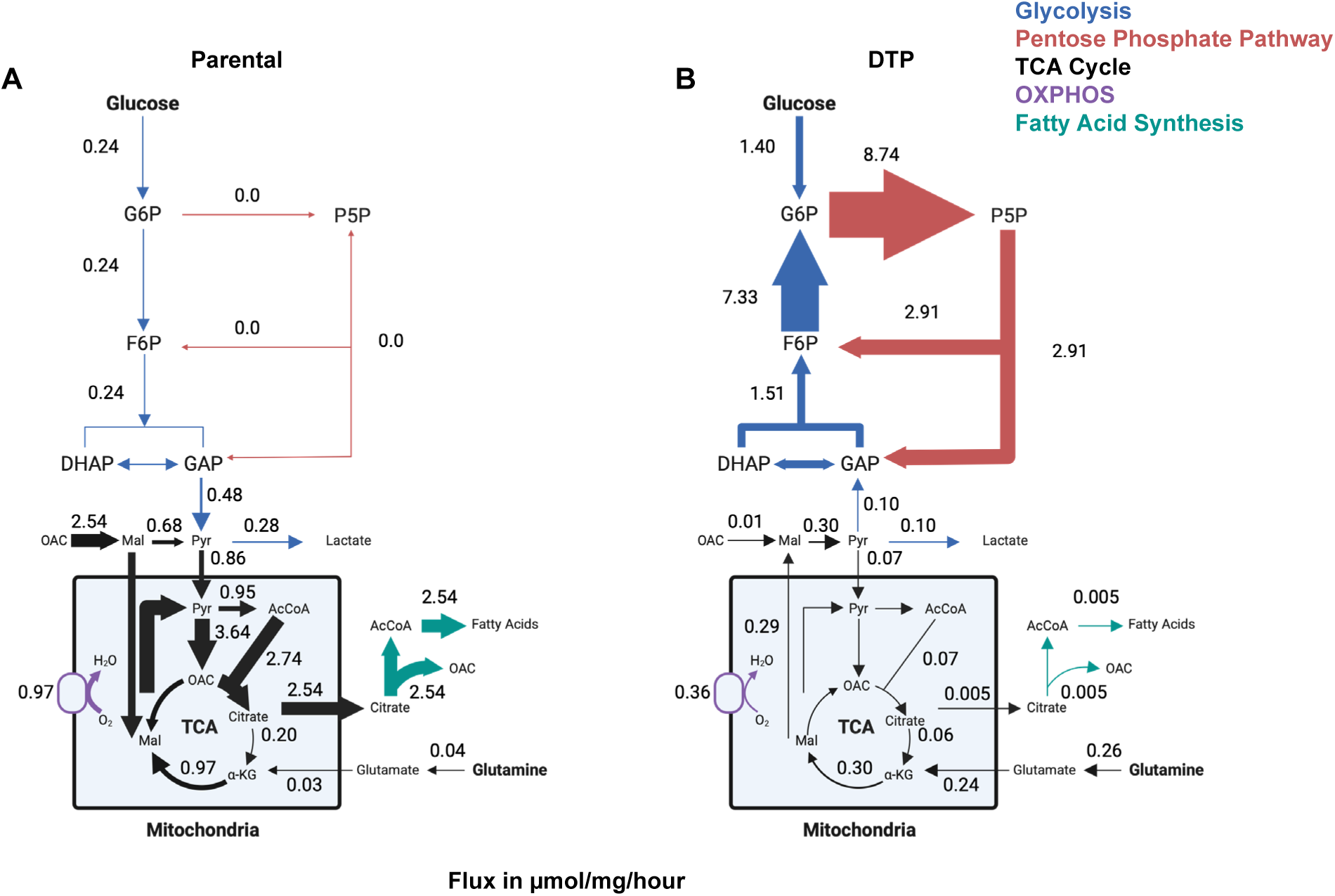
^13^C-metabolic flux analysis reveals the reallocation of glucose and glutamine to fuel the pentose phosphate pathway in DTPs. (**A**) Flux map of central carbon metabolism (in units of µmol per milligram of cell weight per hour) in parental cells. Relative thickness of arrows corresponds to the magnitude of flux. Created in BioRender. Li, M. (2026) https://BioRender.com/r764b3m. (**B**) Flux map of central carbon metabolism (in units of µmol per milligram of cell weight per hour) in DTPs. Relative thickness of arrows corresponds to the magnitude of flux. Created in BioRender. Li, M. (2026) https://BioRender.com/mrphs8q. Numbers are best-fit flux values from ^13^C-metabolic flux analysis (^13^C-MFA) using isotopomer network compartmental analysis (INCA)^29^, constrained by metabolite ^13^C labeling from [U-^13^C] glucose and [U-^13^C] glutamine experiments for both parental cells and DTPs, each with n = 5 biological replicates, and metabolite uptake and excretion rates (n = 3-5 biological replicates).

Mitochondrial metabolism of pyruvate was highly active in parental cells through pyruvate dehydrogenase, generating acetyl-CoA at 0.95 µmol/mg/hr, and pyruvate carboxylase, generating oxaloacetate (OAC) at 3.64 µmol/mg/hr (**Fig. 4A**). Downstream of OAC generation, citrate is synthesized and mostly exported out of the mitochondria in parental cells for fatty acid synthesis, producing cytosolic OAC as a byproduct (**Fig. 4A**). In DTPs, we observed decreased pyruvate import into the mitochondria and a dramatic decrease in citrate export for fatty acid synthesis (**Fig. 4B**). Although DTPs mainly oxidize citrate to alpha-ketoglutarate in the TCA cycle, the flux of that reaction is over 3-fold lower than in parental cells (**Fig. 4A-B**).

Glutamine is another carbon source that feeds into the TCA cycle downstream of citrate. DTPs take up glutamine at a higher rate than parental cells and ultimately synthesize mitochondrial malate in the TCA cycle. Interestingly, DTPs export malate into the cytosol and convert it to pyruvate, which would then feed carbons into the gluconeogenic pathway (**Fig. 4B**). In contrast to DTPs, parental cells retain TCA cycle-derived malate in the mitochondria (**Fig. 4A**). The consumption of TCA cycle metabolites to support gluconeogenesis in DTPs is one possible explanation for the decreased relative pool sizes of intracellular glutamine, glutamate, succinate, and aspartate (**Supplementary Fig. 1A-G**). These data suggest that DTPs metabolize glutamine through the TCA cycle with an anaplerotic function to provide carbons for gluconeogenesis. We hypothesize that the G6P generated from this process and from glucose phosphorylation are then utilized to fuel the oxPPP.

### Reprogramming of central carbon metabolism increases NADPH production and utilization in DTPs

One of the main functions of the oxPPP is to generate a key reducing molecule, NADPH^33^. It can be utilized to reduce oxidized glutathione and replenish reduced thioredoxins for antioxidant defense, consumed for *de novo* fatty acid synthesis, or used by cytochrome P450 (CYP450) enzymes to metabolize xenobiotics^33–35^. Using the quantified metabolic fluxes from our ^13^C-MFA modeling, we estimated NADPH production and consumption fluxes based on known reaction stoichiometry. For example, the flux of NADPH consumption for *de novo* fatty acid synthesis is equal to the fatty acid synthesis carbon flux multiplied by a factor of 1.75, as 14 molecules of NADPH and 8 molecules of acetyl-CoA are consumed to generate one molecule of palmitate^34^. From our central carbon metabolic model, the key reactions that generate NADPH are from the PPP, malic enzyme (ME), glutamate dehydrogenase (GLUD1/2), isocitrate dehydrogenase (IDH1/2), and fatty acid synthesis (**Fig. 5A**). We first calculated the sum of all the fluxes that contributed to NADPH production in each group and subsequently assessed the relative contribution of those reactions to the total NADPH production flux. DTPs exhibited a 3.9-fold increase in total NADPH production based on the reactions described above (**Fig. 5B**). Nearly all of the NADPH produced in PC3 control cells is through cytosolic and mitochondrial malic enzymes (ME1/2), which catalyze the conversion of malate to pyruvate, with a combined flux of 4.14 µmol/mg/hr (**Fig. 5A-B**). On the other hand, nearly all of the NADPH produced in cells 10 Days PTR is through the oxPPP via glucose 6-phosphate dehydrogenase (G6PD) and phosphogluconate dehydrogenase (PGD) at a combined flux of 17.50 µmol/mg/hr (**Fig. 5A-B**). This data indicates that not only is NADPH production increased in DTPs, but that the oxPPP is the dominant contributor of NADPH in DTPs rather than ME1/2 as observed in parental cells. Based on the metabolic steady state assumption and law of mass balance used in ^13^C-MFA, NADPH production must equal NADPH consumption, allowing us to infer that DTPs also exhibit a 3.9-fold increase in total NADPH consumption when compared to parental cells (**Fig. 5B**)^34,36^. Nearly all of the NADPH consumed in parental cells is through fatty acid synthesis, while reactions that were not captured by our core metabolic model, labeled “Other”, contributed the most to NADPH consumption in DTPs (**Fig. 5B**). These reactions include thioredoxin reductase (TXNRD1/2), glutathione reductase (GSR), and peroxiredoxin (PRDX), which are key components in antioxidant defense, and CYP450 (**Fig. 5A**)^37^. Furthermore, DTPs exhibit increased protein expression of NADPH-producing enzymes (G6PD, IDH2, and GLUD2) and antioxidant metabolism enzymes that consume NADPH (**Fig. 5C**). Taken together, these data suggest that DTPs increase NADPH production through the oxPPP and utilize it to fuel antioxidant metabolism.

**Fig. 5.**
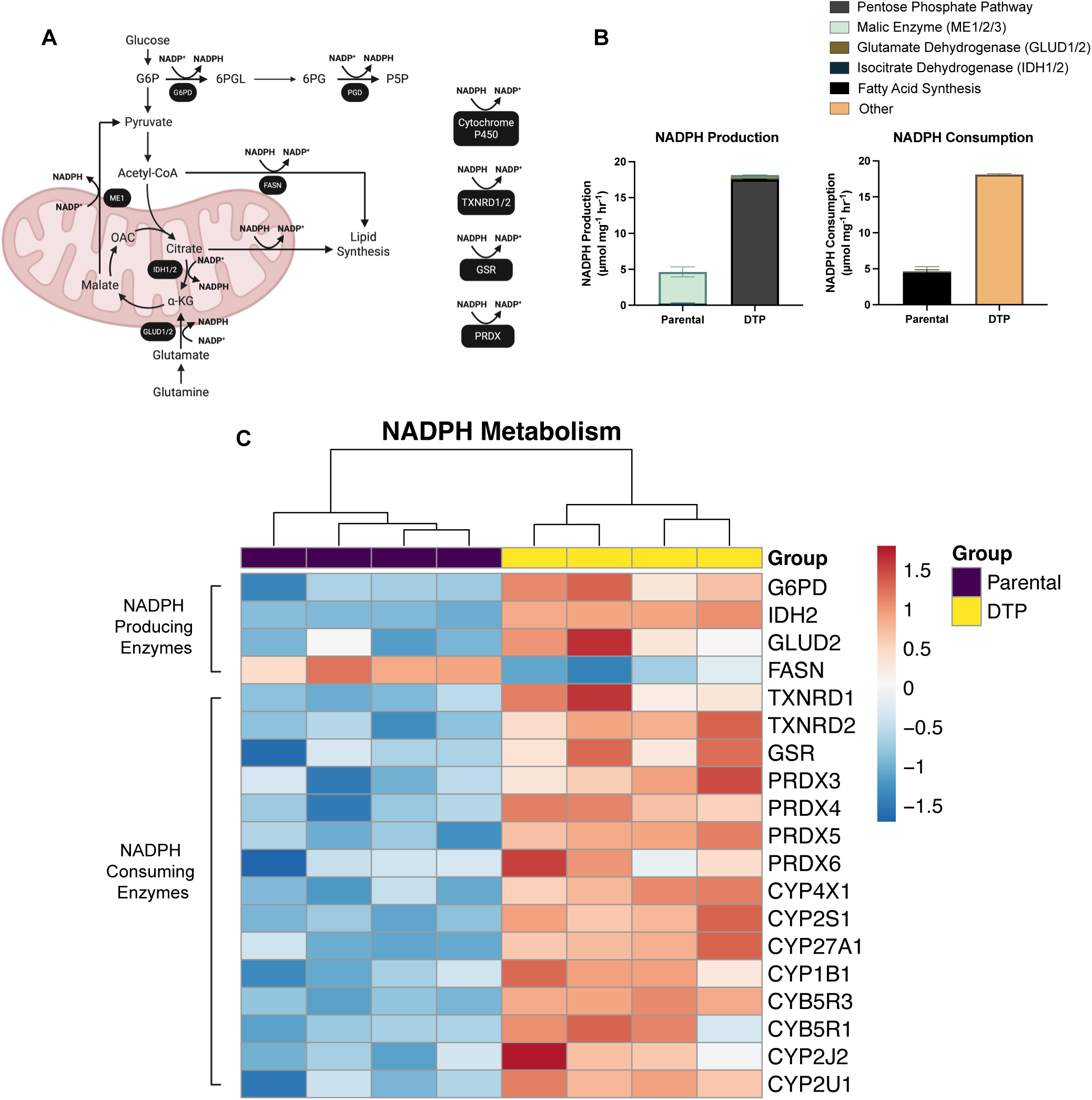
Reprogramming of central carbon metabolism increases NADPH production and utilization in DTPs. (**A**) Schematic of key NADPH-producing and consuming reactions in central carbon metabolism. Enzymes that mediate each reaction are labeled in under the reaction arrow. Created in BioRender. Li, M. (2026) https://BioRender.com/kitiup9. (**B**) NADPH-producing and consuming fluxes in parental cells and DTPs, calculated stoichiometrically from the best-fit flux values from ^13^C-metabolic flux analysis. Relative contribution of pathways to NADPH production and consumption, such as the pentose phosphate pathway, malic enzyme (ME1/2), glutamate dehydrogenase (GLUD1/2), isocitrate dehydrogenase (IDH1/2), fatty acid synthase, and central carbon metabolism-independent reactions, were calculated and compared between parental cells and DTPs. (**C**) Heatmap of differentially expressed proteins that drive NADPH production and consumption in parental cells and DTPs. n = 4 biological replicates (**C**). Data are presented as mean ± s.e.m. (**B**).

### DTPs exhibit an increase in antioxidant metabolism and sustain antioxidant activity following *in silico* knockout of the oxidative pentose phosphate pathway

To assess antioxidant reaction fluxes in DTPs and parental cells, we expanded our metabolic model to the genome scale with the Recon3D metabolic network reconstruction^38^. Our ^13^C-MFA data was limited to only central carbon metabolism, containing 48 essential reactions whereas Recon3D contains 13,543^38^. To better reflect the metabolic state of our cells, the Recon3D model was first reduced to only contain reactions observed in our ^13^C-MFA and bulk proteomics datasets, resulting in a network of 6,039 reactions (**Supplementary Table 1**). We integrated our bulk proteomic expression data and ^13^C-MFA reaction fluxes as constraints in the reduced Recon3D model and performed constraint-based sampling to estimate fluxes of key NADPH-mediated and antioxidant reactions (**Fig. 6A, Supplementary Table 2**). Relative to parental cells, DTPs exhibit increased fluxes of various antioxidant reactions such as cytosolic glutathione oxidoreductase (GTHOr), mitochondrial glutathione oxidoreductase (GTHOm), glutathione peroxidase (GTHPi), and catalase (CATp). In contrast, DTPs reduce flux through thioredoxin reductase (TRDR) (**Fig. 6B**).

**Fig. 6.**
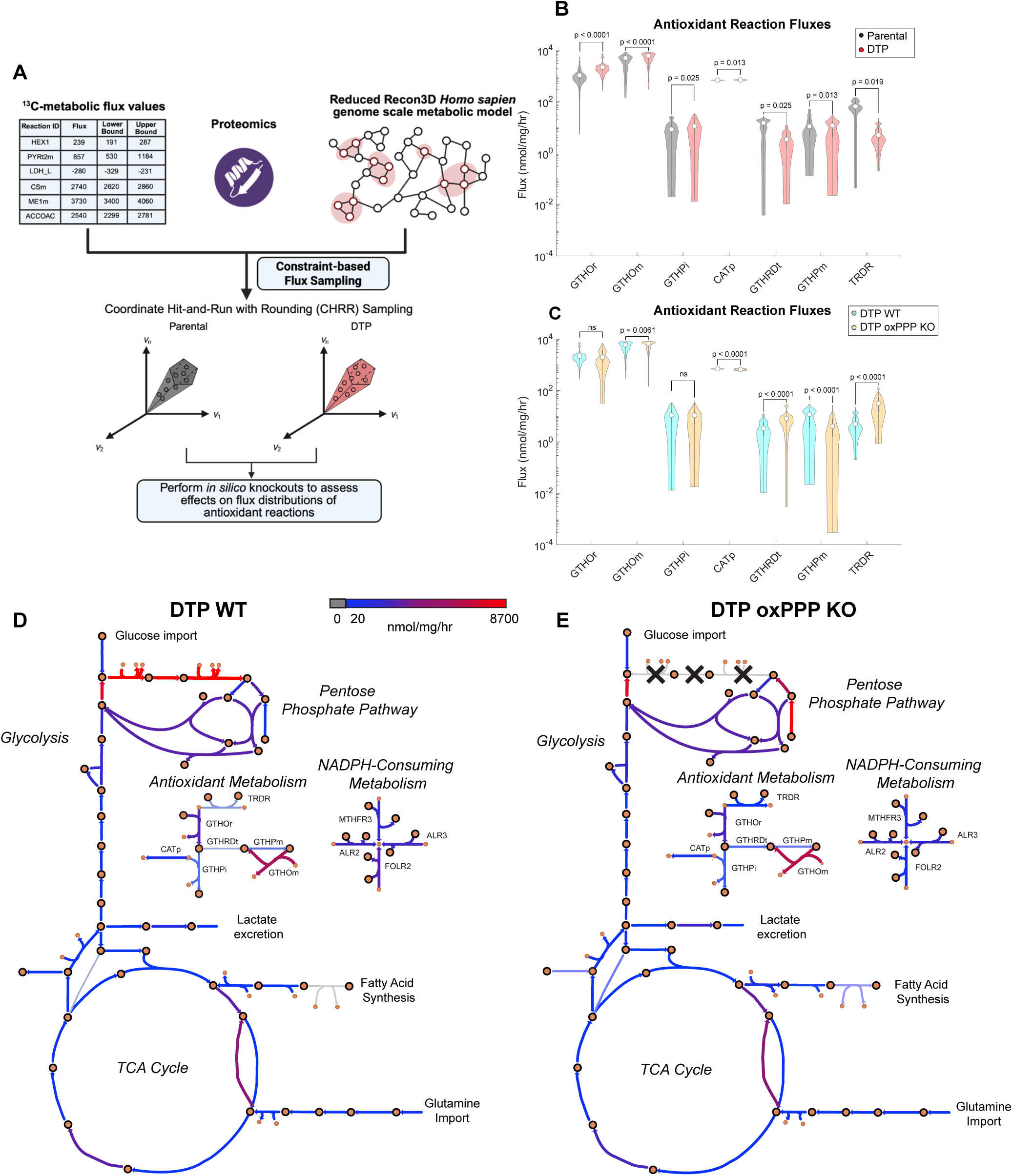
DTPs exhibit an increase in antioxidant metabolism and sustain antioxidant activity following *in silico* knockout of the oxidative pentose phosphate pathway. (**A**) Schematic of genome scale metabolic modeling workflow. Created in BioRender. Li, M. (2026) https://BioRender.com/8fsa6h0. (**B**) Sampled flux estimates of key antioxidant reactions between parental cells and DTPs (in units of nmol/mg/hr). (**C**) Sampled flux estimates of key antioxidant reactions between DTP wild-type (WT) and DTP oxPPP knockout (KO) (in units of nmol/mg/hr). (**D**) Escher map of DTP WT condition. (**E**) Escher map of DTP oxPPP KO condition. Large circles with black outlines represent metabolites, and small circles represent NADPH. Color of arrows correspond to flux value, ranging from 0 to 8700 nmol/mg/hr. n = 1000 CHRR samples per reaction, with 100 skips per CHRR sample (**B-C**). Data are presented as median and interquartile range (**B-C**). P-values were calculated using bootstrapped Mann-Whitney U tests. The median p-value for each reaction is indicated in the figure **(B-C**). Abbreviations: cytosolic glutathione oxidoreductase (GTHOr), mitochondrial glutathione oxidoreductase (GTHOm), cytosolic glutathione peroxidase (GTHPi), catalase (CATp), glutathione transport to mitochondria (GTHRDt), mitochondrial glutathione peroxidase (GTHPm), thioredoxin reductase (TRDR).

One powerful advantage of genome scale metabolic modeling is the ability to perform *in silico* knockouts of specific reactions and assess the effects on other reaction fluxes^39^. We simulated a knockout of the oxPPP in DTPs by setting the G6PD and PGD fluxes to zero and observed an increase in mitochondrial glutathione transport (GTHRDt), mitochondrial glutathione oxidoreductase (GTHOm), and TRDR fluxes, and a decrease in CATp flux (**Fig. 6C**). Interestingly, we observed no changes in GTHOr (**Fig. 6C**). These data suggest that antioxidant metabolism is sustained in response to depletion of oxPPP-derived NADPH. We then hypothesized that DTPs may be altering other NADPH-mediated metabolic reactions in response to oxPPP knockout to balance NADPH pools for antioxidant activity. We observed an increase in NADPH-producing reaction fluxes such as methylenetetrahydrofolate dehydrogenase (MTHFD) and malic enzyme (ME2) in DTP oxPPP knockout when compared to wild-type (**Supplementary Fig. 2A**). Notably, DTPs under oxPPP knockout conditions exhibited decreased flux through NADPH-consuming reactions associated with folate metabolism (**Fig. 6D-E, Supplementary Fig. 2B**). Folate reductase (FOLR2) experienced a 5.40-fold decrease and 5,10-methylenetetrahydrofolate reductase (MTHFR3) experienced a 1.50-fold decrease in flux (**Fig. 6D-E, Supplementary Fig. 2B**). These data suggest that DTPs reduce cytosolic folate metabolism to compensate for the decrease in NADPH production, highlighting their potential metabolic flexibility to retain antioxidant metabolism in response to oxPPP inhibition.

## DISCUSSION

Our study reveals systems-level metabolic shifts that occur in cells that persist after cisplatin treatment. ^13^C-MFA modeling revealed that DTPs increase glucose uptake and shuttle it into the oxPPP (**Fig. 4A-B**). Additionally, DTPs export glutamine-derived carbons out of the TCA cycle to support gluconeogenesis, resulting in the production of G6P that can also fuel the oxPPP (**Fig. 4B**). This data is consistent with the diluted G6P labeling from [U-^13^C] glucose observed in DTPs despite having an increase in glucose consumption rate (**Fig. 2A, D**), as carbons from unlabeled glutamine are utilized anaplerotically to generate G6P via gluconeogenesis. Our differential expression proteomic analysis also revealed that DTPs upregulate key gluconeogenetic enzymes, FBP1 and PCK2 (**Fig. 1C**). Santiappillai et al. reported this phenomenon in a subset of cancer cell lines that shuttled [U-^13^C] glutamine out of the TCA cycle into the gluconeogenic pathway, generating m+3 FBP and phosphoenolpyruvate (PEP)^40^. Others have reported cancer cells that perform glutamine-derived gluconeogenesis increase NADPH production for fatty acid synthesis and other biosynthetic precursors through PCK2 activity^41,42^. In our model, DTPs shuttle glucose- and glutamine-derived carbons to the oxPPP, generating 96% of their intracellular NADPH (**Fig. 5A-B**). Stoichiometric calculations from ^13^C-MFA data suggested that DTPs utilized 99% of the NADPH in metabolic pathways beyond central carbon metabolism, which our model could not resolve (**Fig. 5A-B**).

To address this limitation, we applied genome scale metabolic modeling constrained by empirical metabolic data from DTP and parental cells. Genome scale metabolic modeling is typically performed using flux balance analysis (FBA), which constrains extracellular fluxes and gene or protein expression data to maximize a user-defined cellular objective, such as ATP production or biomass growth^43^. However, the resulting fluxes are predicted by optimizing this objective function and may not accurately represent intracellular fluxes^43^. In this study, we integrated flux estimates from ^13^C-MFA based on data from isotope tracing of intracellular nutrient utilization into the genome scale model constraints to more accurately reflect the metabolic state of our cells. In addition, we performed flux sampling through CHRR-MCMC rather than optimizing for a singular cellular objective, allowing us to assess metabolic flux distributions at the genome scale in DTPs and parental cells with 95% confidence intervals^44^.

Genome scale metabolic modeling revealed that DTPs increase antioxidant reaction fluxes when compared to parental cells (**Fig. 6B**). These data align with previous work from our group showing DTPs surviving cisplatin chemotherapy exhibit increased levels of reactive oxygen species (ROS) and enriched NRF2-mediated antioxidant response when compared to parental cells^45,46^. Furthermore, we previously identified that inhibiting glutathione peroxidase 4 (GPX4) selectively induced ferroptosis in DTPs, which has also been reported by other groups^45,47,48^. Metabolic CRISPR screens have also revealed that NRF2-driven antioxidant metabolism and the pentose phosphate pathway are required for DTP survival^49^. Taken together, these findings demonstrate that antioxidant metabolism plays a critical role for survival in the DTP state.

When testing the essentiality of the oxPPP for antioxidant metabolism in DTPs *in silico*, we identified changes in other NADPH-producing and consuming fluxes while detecting minimal loss of antioxidant fluxes. Notably, we observed elevated mitochondrial NADPH production via MTHFD in response to simulated oxPPP knockout, which could explain the increase in mitochondrial glutathione antioxidant activity through GTHRDt and GTHOm (**Fig. 6C, Supplementary Fig. 2A**). DTPs under oxPPP knockout also exhibited increased flux through malic enzyme (ME2) when compared to wild type DTPs (**Supplementary Fig. 2A**). Our findings are consistent with previous studies that reported cancer cells with a loss-of-function in G6PD increase flux through ME as a compensatory mechanism to generate NADPH^50,51^.

Concurrently, we also observed decreases in NADPH-consuming fluxes related to folate metabolism – FOLR2 and MTHFR3 (**Fig. 6D-E; Supplementary Fig. 2B)**. We hypothesize that DTPs decrease folate metabolism as a compensatory mechanism to sustain intracellular NADPH pools for antioxidant activity. This is consistent with observations of folate deficiency and impaired dihydrofolate reductase (DHFR) activity in response to G6PD knockout across cancer cell lines^50^. As DHFR is the main enzyme that facilitates the FOLR2 reaction and generates tetrahydrofolate (THF), a precursor for MTHFD^38^, we hypothesize that further inhibiting DHFR may induce depletion of THF that feeds into MTHFD, mitigating the elevated NADPH production we observed in DTPs under oxPPP knockout. Thus, residual DHFR activity may remain essential for survival in the DTP state.

These findings suggest a potential synthetic lethality treatment paradigm involving co-treatment of G6PD inhibitors, such as 6-aminonicotinamide, and DHFR inhibitors, such as methotrexate, to eliminate DTPs surviving cisplatin chemotherapy^52,53^. Future work involves testing this hypothesis with cell viability experiments to assess synergism between G6PD and DHFR inhibition on DTPs. In summary, we propose that DTPs exhibit a systems-level shift in central carbon metabolism to sustain antioxidant activity for cell survival (**Fig. 7**). Additional characterization of the adaptive metabolic responses of the DTP state with integrated fluxomics and functional validation will reveal new vulnerabilities that can be leveraged for targeting and improve patient prognosis.

**Fig. 7.**
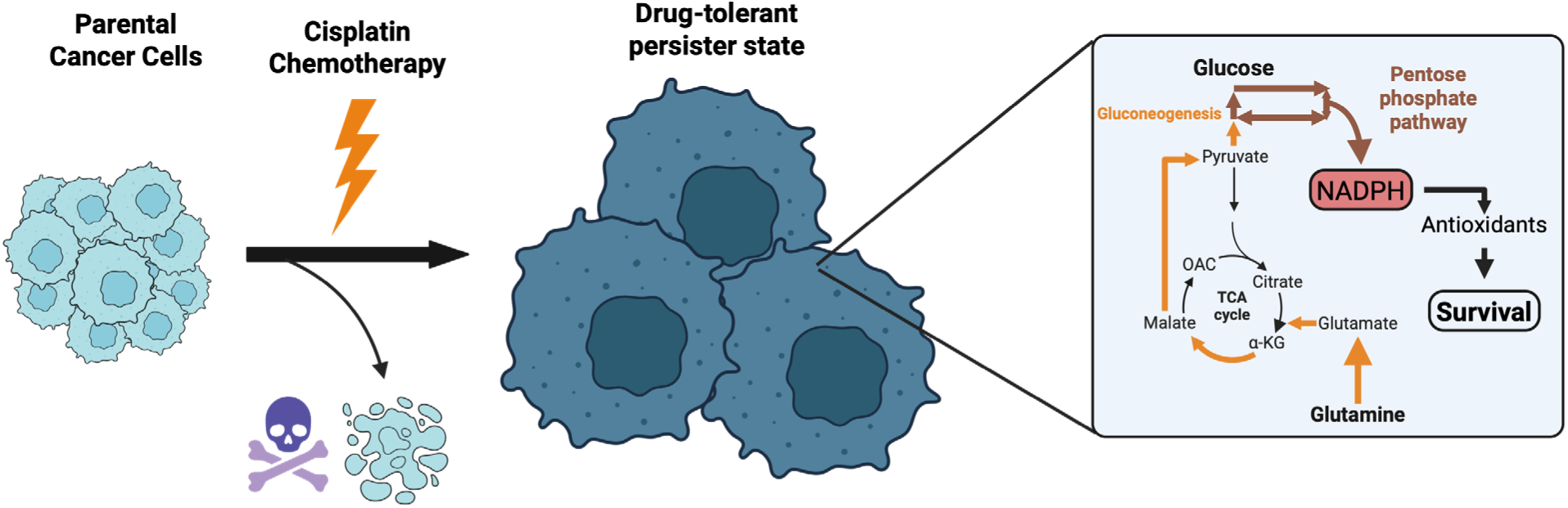
DTPs reallocate central carbon sources to sustain antioxidant metabolism for cell survival. Cancer cells that survive cisplatin chemotherapy enter a drug-tolerant persister (DTP) state, presenting with enlarged cell size and proliferative arrest. Cells in this DTP state shuttle glucose into the pentose phosphate pathway, generating NADPH. Carbons derived from glutamine are metabolized in the TCA cycle and subsequently exported out of the mitochondria to fuel gluconeogenesis, ultimately feeding into the pentose phosphate pathway. The NADPH generated through these processes are consumed by antioxidant reactions, which are essential for survival in the DTP state. This proposed mechanism requires further validation with future studies. Created in BioRender. Li, M. (2026) https://BioRender.com/x1g1040.

## MATERIALS AND METHODS

### Cell culture and induction of DTP cell state

PC3 parental cells were cultured in RPMI 1640 medium (ThermoScientific; Cat# 11875119) with 10% fetal bovine serum (FBS) (Avantar; Cat# 97068-085) and 1% penicillin and streptomycin (P/S) (Gibco; Cat#15140-122). Cells were incubated at 37℃ and 5% CO_2_. To induce the DTP cell state, cells were treated with the LD_50_ dose of cisplatin (6µM) for 72 hours. Drug-containing media was removed, and cells were left in culture for 10 days, with media changes every 3 days. Cells at 10 days post-treatment removal (DTPs) and parental cells were used for experiments.

### Protein lysate harvest for bulk proteomics

Parental cells and DTPs were lifted from T150 flasks with TrypLE^TM^ Express (GibcoTM; Ref# 12604-013) and lysed using RIPA buffer (Sigma-Aldrich; Cat#: R0278-500ML) with 1X Halt Protease & Phosphatase cocktail and 0.005 M EDTA (ThermoScientific; Cat#: 78440). Lysates were sonicated in a Branson 3510 Ultrasonic Cleaner for 10 minutes and then were subsequently spun in a 4℃ centrifuge at 21,000 rcf. Supernatant was collected and stored at - 20℃. The Pierce^TM^ BCA Protein Assay (ThermoScientific; Cat#: 23227) was used to quantify protein concentration, and 10 µg of protein was used for bulk proteomics.

### Sample preparation for bulk proteomics

Protein extracts were buffer exchanged using SP3 paramagnetic beads (GE Healthcare)^54^. Briefly, 10 µg of protein was resolubilized in 10 mM TEAB and disulfide bonds reduced with dithiothreitol (5 mM final concentration) for 1 hour at 60℃. Samples were cooled to RT and pH adjusted to ∼8.0, followed by alkylation with iodoacetamide (10 mM final concentration) in the dark at RT for 15 minutes. Next, SP3 beads were added to the samples at a ratio of 10:1 bead:protein and 100% ethanol was added to achieve a final ethanol concentration of 50% v/v. Samples were incubated at RT with shaking for 5 minutes. Following protein binding, beads were washed with 180 µL 80% ethanol three times. Proteins were digested on-bead with trypsin (Pierce) at 37℃ overnight at a ratio of 10:1 protein:enzyme. Resulting peptides were separated from the beads using a magnetic tube holder. Supernatants containing peptides were removed from the beads and dried using vacuum centrifugation.

### Isobaric tandem mass tag (TMT) labeling

Peptides were labeled with TMTpro reagents (Thermo Fisher) according to the manufacturer’s instructions. After quenching of the labeling reaction with hydroxylamine (0.2% v/v final concentration), labeled peptide samples were combined and dried by vacuum centrifugation.

### Peptide fractionation

The combined TMT-labeled peptides were re-constituted in 100 µL 100mM TEAB buffer and filtered through EasyPep Mini cleanup columns (Thermo Fisher Scientific) to remove excess TMT label and hydroxylamine. Peptides in the flow through were diluted to 2 mL in 10 mM TEAB in water and loaded on a XBridge C18 Guard Column (5 µm, 2.1 × 10 mm, Waters) at 250 µL/min for 8 min prior to fractionation on a XBridge C18 Column (5 µm, 2.1 × 100 mm column (Waters) using a 0 to 90% acetonitrile in 10 mM TEAB gradient over 85 min at 250 µL/min on an Agilent 1200 series capillary HPLC with a micro-fraction collector. Eighty-four 250 µl fractions were collected and concatenated into 24 fractions according to the method described by Wang et al. and dried^55^.

### Liquid chromatography separation and tandem mass spectrometry (LC-MS/MS) for proteomics

Dried peptides were reconstituted in 2% ACN and 0.1% FA and analyzed by nanoflow liquid chromatography-tandem mass spectrometry (nLC-MS/MS) using a Neo Vanquish UHPLC interfaced with an Orbitrap Exploris 480 mass spectrometer (both instruments, Thermo Fisher Scientific). Peptide separation was performed with linear gradients (water/acetonitrile) over polyimide-coated, fused-silica, 25 cm × 360 μm o.d./75 μm i.d. self-packed columns with Kasil frits and PepSep stainless steel emitter (30 µm inner diameter, Bruker Daltonics). Stationary phase in analytical columns consisted of ReproSil-Pur 120 C18-AQ, 2.4 μm particle size, 120 Å pore (Dr. Maisch High Performance LC GmbH). Trap columns consisted of ∼1 cm × 360 μm o.d./75 μm i.d. polyimide-coated, fused-silica tubing (New Objective), packed with 5 μm particle size, 120 Å pore, C18 stationary phase (ReproSil-Pur), with a Kasil frit. Electrospray ionization was accomplished with 2 kV positive spray voltage and an ion transfer tube temperature of 250 °C. For DDA TMT analysis, each peptide fraction was separated by a 120-minute linear gradient. MS1 precursor ion scans were acquired in the Orbitrap detector of the Exploris 480 mass spectrometer with a 3 second cycle time from 400-1500 m/z at 120,000 resolution at 200 m/z with a normalized AGC of 300%, an RF lens setting of 50%, and maximum injection time set to Auto. Precursor ions were individually isolated with a 0.7 m/z isolation window and a 3-second duty cycle, with the following filters: monoisotopic precursor selection (MIPS) set to peptide mode, intensity threshold 2.5×104, charge states 2-6, dynamic exclusion duration of 45 s (10 ppm tolerance), isolation purity 70%. Precursors were fragmented by HCD with a 36% normalized collision energy and MS/MS spectra were acquired at 30,000 resolution, AGC set to Standard and maximum injection time set to Auto.

### Bulk proteomics data analysis

All database searches were done in Proteome Discoverer v3.1 (ThermoFisher Scientific) with Chimerys using the Inferys v3.0 prediction model. For TMT analysis, all fractions were searched together against a human UniProt FASTA database (proteome accession UP000005640, 98758 entries) and a custom database containing common contaminants (e.g., human keratin; 438 entries). Search criteria were tryptic cleavage (maximum 2 missed), peptide length 7-30, peptide charge 2-6, 10 ppm fragment ion mass tolerance, Cys carbamidomethylation, Lys TMTpro labeling, and N-terminal TMTpro labeling as fixed modifications, and Met oxidation as a variable modification (max 3/peptide). Peptide identifications were validated by Chimerys at 1% false discovery rate (FDR) based on an auto-concatenated decoy database search. Peptide spectral matches (PSMs) were filtered for isolation interference ≤30% and a normalized Chimerys coefficient of 0.8. Only proteotypically unique peptides were used for relative quantitation. TMT reporter ions were normalized to total peptide amount per channel and quantification on reporter ion signal-to-noise values. Group differences were tested by ANOVA based on protein abundance with no imputation of missing values. Differential protein expression analysis, principal component analysis, gene set enrichment analysis, and plots were generated in RStudio with the following packages: limma, FactoMineR, EnhancedVolcano, clusterProfiler, factoextra, enrichplot, and ggplot2 ^56–61^. Data used for analysis are included in **Source Data Fig. 1.**

### Extracellular flux measurements of metabolite concentrations in media

During the [U-^13^C] glucose and [U-^13^C] glutamine tracing experiments, 300 µL of media was obtained for analysis every hour, flash frozen, and stored at −80 ℃. For glucose, lactate, glutamine, and glutamate concentration measurements, media were input into a YSI 2900 Biochemistry Analyzer (YSI) equipped with Glucose/Lactate and Glutamine/Glutamate modules. Metabolite concentrations were converted to metabolite abundance and normalized to cell weight. The normalized metabolite abundances were used to determine the rate of change over time via linear regression analysis. The extracellular fluxes were plotted as nmol per milligram of cell weight per hour.

### Isotope tracing with [U-^13^C] glucose, [1,2-^13^C] glucose, and [U-^13^C] glutamine

Cells were cultured to the desired timepoints. On the day of the experiment, [U-^13^C] glucose (Cambridge Isotope Laboratories; Cat#: CLM-1396-10) or [1,2-^13^C] glucose (Cambridge Isotope Laboratories; Cat#: CLM-504-1) was added to RPMI 1640 Glucose-Free Media (ThermoScientific; Cat# 11879020) at a final concentration of 11.11 mM with 10% dialyzed FBS (ThermoScientific; Cat#: 26400044), and 1% Penicillin/Streptomycin (Gibco; Cat#15140-122). Cells were incubated with [U-^13^C] glucose or [1,2-^13^C] glucose tracer for 3 hours and 6 hours. At endpoint, cells were lifted with TrypLE^TM^ Express (Gibco^TM^; Ref# 12604-013), which was subsequently neutralized with dialyzed FBS-containing media and spun down at 500g for 5 minutes at 4 ℃. Cells were washed with 1X PBS twice and pelleted into a 1.5 mL Eppendorf tube. Cell weights were measured, and pellets were snap frozen with liquid nitrogen and stored at −80℃ until metabolite extraction. For [U-^13^C] glutamine experiments, [U-^13^C] glutamine (Cambridge Isotope Laboratories; Cat#: CLM-1822-0.5) was added to RPMI 1640 Glutamine-Free Media (ThermoScientific; Cat# 11879020) at a final concentration of 2.20 mM with 10% dialyzed FBS (ThermoScientific; Cat#: 26400044), and 1% Penicillin/Streptomycin (Gibco; Cat#15140-122). Cells were incubated with [U-^13^C] glutamine tracer for 3 hours and 6 hours. At endpoint, cells were lifted with TrypLE^TM^ Express (Gibco^TM^; Ref# 12604-013), which was subsequently neutralized with dialyzed FBS-containing media and spun down at 500g for 5 minutes at 4 ℃. Cells were washed with 1X PBS twice and pelleted into a 1.5 mL Eppendorf tube. Cell weights were measured, and pellets were snap frozen with liquid nitrogen and stored at −80℃ until metabolite extraction. Cells underwent freeze thaw cycles in liquid nitrogen followed by homogenization in 1:1 methanol:water using needle sonication. The samples were then centrifuged at 5,000 RPM for 10 minutes at 4 ℃. Samples were filtered using a 3 kDa Amicon filter to remove proteins and lipids, dried using a speed vacuum, and then reconstitutied in a 1:1 methanol:water mixture. Glycolytic intermediates and tricarboxylic acid (TCA) cycle metabolites and their isotopomers were quantified using a Luna NH2 column (3 µm, 150 x 2 mm, Phenomenex, Torrance, CA), following previously described detailed methods^62–65^. Natural abundance correction was performed with IsocorrectoR^66^ and mass isotopomer distributions (MIDs) were plotted in GraphPad Prism as “% of pool”, indicating the relative amount of a specific mass isotopomer divided by the sum of all isotopomers of the metabolite and multiplied by 100. Data plotted in GraphPad Prism are included in **Source Data Fig. 2 and Source Data Fig. 3.**

### Quantification of [1,2-^13^C] glucose contribution to lactate

Sample preparation was performed based on a protocol from Ahn & Antoniewicz^67^. Briefly, media samples were dried under nitrogen gas flow at 37 ℃. The dried samples were then resuspended in 2% methoxylamine hydrochloride in pyridine and subsequently heated at 37 ℃. Derivatization was performed with the addition of N-methyl-N-(tert-butyldimethylsiyl)-trifluoroacetamide (MTBSTFA) + 1% tertbutyldimethylchlorosilane (TBDMCS) and incubation at 60 ℃. Samples were spun down at max speed for 5 minutes and supernatant was transferred to gas chromatography vials for GC-MS. GC-MS was performed using the Agilent 7890A GC connected to an Agilent 5977B Mass Spectrometer. Peak calling and integration were performed using PIRAMID^68^.

### ^13^C-metabolic flux analysis (^13^C-MFA)

Metabolic flux analysis was performed with the INCA 2.4 software based on the elementary metabolite unit (EMU) framework^29,36^. Mass isotopomer distributions (MIDs) of measured metabolites from [U-^13^C] glucose and [U-^13^C] glutamine tracer experiments and measured extracellular fluxes (glucose and glutamine import, lactate export, and oxygen consumption) were integrated into a defined isotopomer network model of central carbon metabolism. The oxygen consumption flux was determined previously by our group^46^. Fluxes were estimated by minimizing the sum of squared residuals (SSR) between simulated and observed MIDs^29,36^. Best-fit flux values were determined through the least squares regression of 100 random initial guesses. A chi-squared test was performed to assess goodness-of-fit of the flux solutions, and parameter continuation was performed to obtain 95% confidence intervals for each calculated flux^29^. Model fits for both parental cells and DTPs were accepted with SSRs within their respective 95% confidence intervals. Raw data outputs from INCA 2.4 are included in Source Data Fig. 4.

### Genome scale metabolic modeling

Genome scale metabolic modeling using the the Recon3D metabolic network was performed through Constraint-Based Optimization and Reconstruction Analysis (COBRA) in MATLAB (MathWorks)^38,39^. Proteomic integration into the model was performed based on the E-Flux method, where empirical protein expression data was used to set upper and lower bounds for reaction fluxes based on the assumption that higher protein expression leads to higher flux^69^. log2-transformed proteomics expression data was globally normalized so that the highest expression value was equal to 1. These normalized expression values were used as coefficients to set flux bounds for their corresponding reactions. The default maximum flux for each reaction was set to 10,000 nmol/mg/hr. The maximum flux of a certain reaction after proteomic integration was the default value multiplied by the expression coefficient. Flux data from ^13^C-MFA were used to set constraints on corresponding reactions in the reduced Recon3D model. No objective function was set. Coordinate hit-and-run with rounding Markov-Chain Monte Carlo (CHRR-MCMC) analysis was performed to sample flux distributions from the constrained metabolic models of DTP and parental cells^44^. For each parental and DTP model, the flux space was defined based on the proteomic and ^13^C-MFA constraints. Flux sampling was performed on 1000 samples, where the number of skips between samples was set to 100. P-values were calculated using bootstrapped Mann-Whitney U tests. For each of the 100 bootstrap iterations, 100 samples were randomly selected from each group, and a Mann-Whitney U test was performed. The median p-value for each reaction is indicated in the figure. Raw data output from CHRR-MCMC sampling are included in **Source Data Fig. 6.** Metabolic maps were visualized in Escher^70^. Escher maps and corresponding data input from CHRR-MCMC sampling are included in **Supplementary Data 1** to visualize metabolic fluxes at https://escher.github.io/. Nomenclature of all the reactions from genome scale modeling is from the Recon3D model reconstruction^38^.

### Statistics

Statistical analyses were performed in GraphPad Prism version 9.1 (GraphPad Software, LLC) and R. Statistical tests that were performed, and p-values are reported in respective figure legends. An *α* value of 0.05 was used for all tests.

## DATA AVAILABILITY

Data is provided within the manuscript or in supplementary information files. Source data for each figure is provided in Microsoft Excel format. Code used to perform genome scale metabolic modeling is available at Github (https://github.com/mli154/genome-scale-modeling-with-proteomics-and-13C-MFA).

## ACKNOWLEDGEMENTS

We thank the William and Carolyn Stutt Research Fund, Ronald Rose, MC Dean, Inc., William and Marjorie Springer, Mary and Dave Stevens, Louis Dorfman, and the Jones Family Foundation for their support of our work. We thank the Center for Proteomics Discovery at the Johns Hopkins School of Medicine for the generation of the bulk proteomics dataset. We thank the Baylor College of Medicine Metabolomics Core for mass spectrometry data generation for the intracellular isotope tracing experiments. Work performed by the Baylor College of Medicine Metabolomics Core was supported by CPRIT Proteomics and Metabolomics Facility (RP210227), NIH (P30CA125123), and Dan L. Duncan Cancer Center.

## AUTHOR CONTRIBUTIONS

Conceptualization: ML, KJP, SRA; Experimental work: ML, BP, LVL; Data analysis and figure generation: ML, BP; Writing – original draft preparation: ML; Writing – review and editing: ML, KJP, SRA. Funding acquisition: KJP, SRA, MJB.

## ETHICS DECLARATIONS

### Approval for animal experiments

Not applicable

### Approval for human experiments

Not applicable

### Consent to participate/consent to publish

Not applicable

