## Supplementary Information for "Drug-tolerant persister cells reallocate carbon sources to fuel antioxidant metabolism for survival"

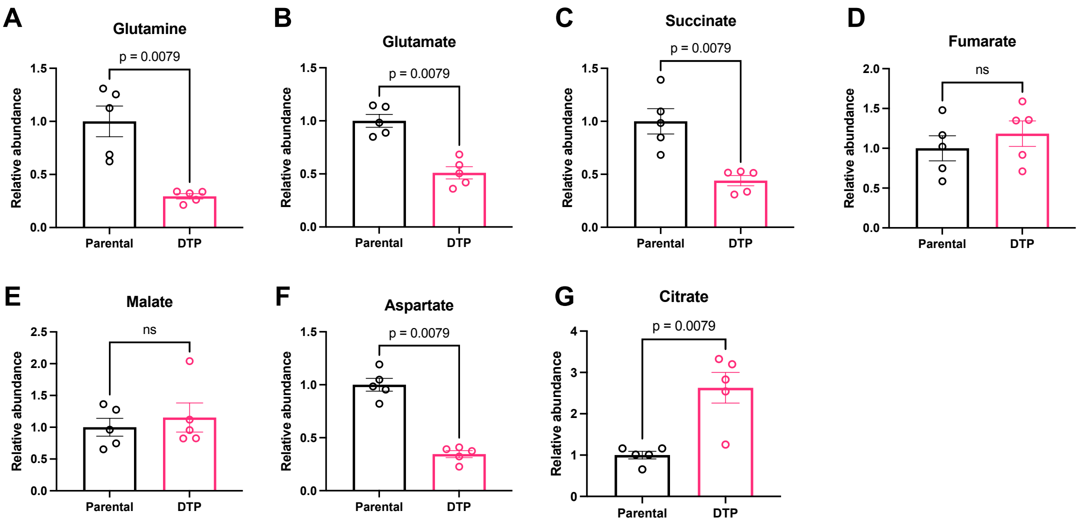


**Supplementary Fig. S1**. **Relative pool sizes of TCA cycle metabolites in parental cells and DTPs**. (**A**) Glutamine. (**B**) Glutamate. (**C**) Succinate. (**D**) Fumarate. (**E**) Malate. (**F**) Aspartate. (**G**) Citrate. n = 5 biological replicates for parental cells and DTPs. Data are presented as mean ± s.e.m. P-values were calculated using a two-tailed Mann-Whitney U test and exact values are indicated in the figure.

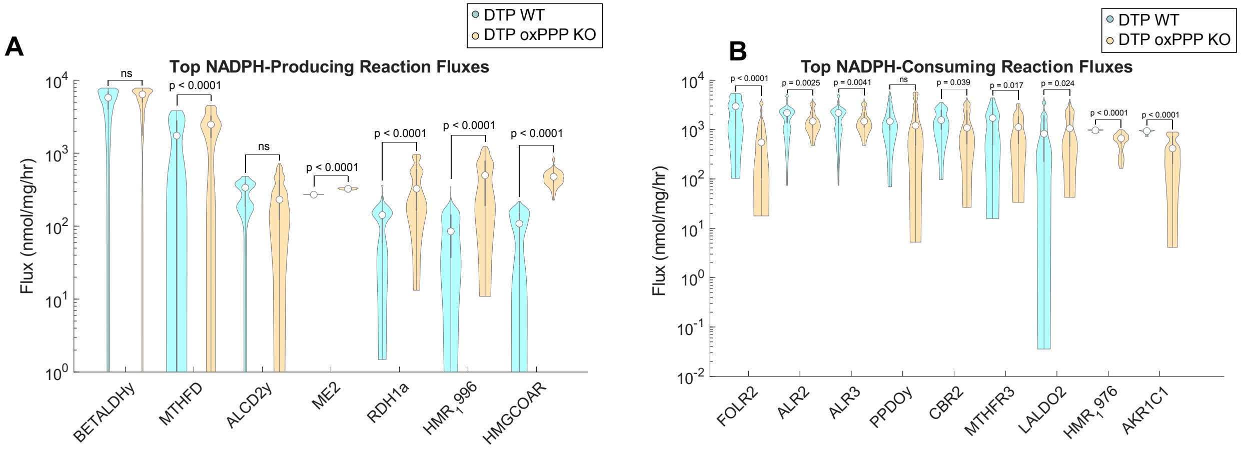


**Supplementary Fig. S2**. **DTPs exhibit increased NADPH production and decreased NADPH consumption upon knockout of the oxidative pentose phosphate pathway**. (**A**) Sampled flux estimates of key NADPH-producing reactions between DTP wild-type (WT) and DTP oxPPP knockout (KO) (in units of nmol/mg/hr). (**B**) Sampled flux estimates of key NADPH-consuming reactions between DTP wild-type (WT) and DTP oxPPP knockout (KO) (in units of nmol/mg/hr). n = 1000 CHRR samples per reaction, with 100 skips per CHRR sample. Data are presented as median and interquartile range. P-values were calculated using bootstrapped Mann-Whitney U tests. For each of the 100 bootstrap iterations, 100 samples were randomly selected from DTP WT and KO groups, and a Mann-Whitney U test was performed. The median p-value for each reaction is indicated in the figure. Abbreviations: betaine-aldehyde dehydrogenase (BETALDHy), methylenetetrahydrofolate dehydrogenase (MTHFD), alcohol dehydrogenase (ALCD2y), malic enzyme (ME2), retinol dehydrogenase (RDH1a), 11Beta-hydroxysteroid dehydrogenase (HMR_1996), hydroxymethylglutaryl CoA reductase (HMGCOAR), folate reductase (FOLR2), aldose reductase (ALR2/3), propane-1,2-diol:NADP+ 1-oxidoreductase (PPDOy), carbonyl reductase (CBR2), 5,10-methylenetetrahydrofolate reductase (MTHFR3), D-lactaldeyhyde:NADP+ 1-oxidoreductase (LALDO2), 3-Oxo-5Alpha-steroid 4-dehydrogenase (HMR_1976), aldo-keto reductase family 1, member C1 (AKR1C1).

**Description of Additional Supplementary Files**

**File Name:** Supplementary Table 1
**Description**: Reduced Recon3D metabolic network reconstruction used for genome scale metabolic modeling. Number of reactions were reduced from 10,543 to 6,039 based on proteomic and $^{13}C$-metabolic flux analysis data. This table consists of reaction identifier in Recon3D nomenclature, the reaction name, and the reaction formula.

**File Name**: Supplementary Table 2
**Description**: Key antioxidant and NADPH-mediated reactions assessed from genome scale metabolic modeling between parental cells, DTP wild-type, and DTP oxPPP KO. This table consists of reaction identifier in Recon3D nomenclature, the reaction name, and the reaction formula.

**File Name**: Supplementary Data 1
**Description**: Escher map .json file with input data of DTP wild-type and oxidative pentose phosphate pathway knockout fluxes for visualization at https://escher.github.io/.

**File Name**: Source Data Fig. 1
**Description**: Source data for Figure 1.

**File Name**: Source Data Fig. 2
**Description**: Source data for Figure 2.

**File Name**: Source Data Fig. 3
**Description**: Source data for Figure 3.

**File Name**: Source Data Fig. 4
**Description**: Source data for Figure 4.

**File Name**: Source Data Fig. 5
**Description**: Source data for Figure 5.

**File Name**: Source Data Fig. 6
**Description**: Source data for Figure 6.

**File Name**: Source Data Supplementary Fig. S1
**Description**: Source data for Supplementary Fig. S1.

**File Name**: Source Data Supplementary Fig. S2
**Description**: Source data for Supplementary Fig. S2.
